# A latitudinal gradient in the diel partitioning of species richness?

**DOI:** 10.1101/2024.05.20.594953

**Authors:** Mark K. L. Wong

## Abstract

The writings of naturalists from two centuries past are brimming with accounts of the stark differences in the kinds and numbers of organisms encountered during the day and night and between the tropical and temperate zones. However, only recently have ecologists begun to systematically describe and explain the geographic variation in the diel activities of species on Earth. Examining data from 60 insect communities globally, I find that the partitioning of total species richness across three diel activity periods tracks the latitudinal gradient. In general, the proportions of diurnal and nocturnal species are highest among tropical communities and decline poleward, while cathemeral activity characterises over half of all species in communities at high latitudes. These latitudinal trends in diel partitioning at the community level broadly reflect recently documented patterns in the global distributions of vertebrate species using different activity periods. I outline six hypotheses that may account for a latitudinal gradient in the diel partitioning of species richness.

## Introduction

> “*The moths in certain families, such as the Zygaenidae, various Sphingidae, Uraniidae, some Arctiidae and Saturniidae, fly about during the day or early evening, and many of these are extremely beautiful, being far more brightly coloured than the strictly nocturnal kinds*.”

– Charles Darwin, 1871, *The Descent of Man*

The contrasting identities and traits of animals active during the day and night were certainly not lost on the early naturalists. Ecological studies have since revealed that the partitioning of diel activity time among such diurnal and nocturnal species in a community can serve as an important coexistence mechanism (Carothers & Jaksić, 1984; Kronfeld-Schor & Dayan, 2003; Lear et al., 2021). Today, there is growing interest in the diel variation of biodiversity-mediated ecosystem functions such as pollination, herbivory, and nutrient cycling (Cox & Gaston, 2024). Furthermore, there is widespread concern that specific diel communities are disproportionately susceptible to the varying levels of anthropogenic light, noise, and chemicals in environments over the 24-h diel cycle (Gaynor et al., 2018; Sanders et al., 2021; Thoré et al., 2024).

Despite growing appreciation for the diel dynamics of biodiversity, as well as mounting evidence that cathemerality (activity during both day and night) is a distinct and prevalent diel strategy across the animal kingdom (Tattersall, 1987; Cox & Gaston, 2024), there remains a dearth of fundamental information on the relative proportions of diurnal, nocturnal, and cathemeral species within communities. This problem – symptomatic of the diurnal habits of ecologists (Gaston, 2019) – is arguably most severe for insects (Wong & Didham, 2024), a functionally diverse and increasingly imperilled group (Janzen & Hallwachs, 2021).

> “*The nearer we approach the tropics, the greater the increase in the variety of structure, grace of form, and mixture of colours, as also in perpetual youth and vigour of organic life*.”

– Alexander von Humboldt, 1807, *Views of Nature* (1850 translation)

Beyond noting changes in animal activity over the diel cycle, pioneering naturalists were captivated by the staggering increments in the numbers of species they encountered while journeying from temperate zones toward the tropics. Following in their footsteps, modern ecologists have sought to systematically document and explain latitudinal patterns in species richness (Pianka, 1966; Willig et al., 2003) and more recently in functional diversity (Schumm et al., 2019), genetic diversity (Miraldo et al., 2016), and biotic interactions (Roslin et al., 2017).

Given that key factors influencing diel activity such as temperature and light also vary with latitude (Bennie et al., 2014), the relative proportions of species using different diel periods in a locality may likewise depend on its distance from the equator. Indeed, recent efforts to map the global distributions of diurnal, nocturnal, and cathemeral mammals (>4,400 species, Bennie et al., 2014; Cox & Gaston, 2024) and lizards (>3500 species, Vidan et al., 2019) revealed broad latitudinal trends. They found that the richness of both nocturnal and diurnal species peaked in the tropics and declined poleward, while that of cathemeral species was highest at mid-to-high latitudes (∼40°–70°) (Bennie et al., 2014; Vidan et al., 2019; Cox & Gaston, 2024). Notably, these patterns were documented based on the distributions of vertebrate species across geographic regions globally. Therefore, it remains to be seen whether latitudinal gradients likewise shape the diel partitioning of species richness at the finer scale of the ecological community, and particularly among invertebrate species.

To survey diel patterns in insect community structure, I recently searched the literature for studies which had systematically sampled insect communities in comparable day and night periods using methods that exclusively targeted active insects. Multiple studies identified during the search did not report sample-based estimates of species richness and had to be excluded from a subsequent meta-analysis, which mainly examined diel patterns in insect abundance (Wong & Didham, 2024). Nonetheless, many studies reported the total numbers of species collected across all samples – and crucially – the numbers collected exclusively in day samples (diurnal species), exclusively in night samples (nocturnal species), and in both day and night samples (cathemeral species). Here I summarise this valuable information and present evidence of a latitudinal gradient in the diel partitioning of species richness in insect communities globally. The goal of this paper is to ignite serious contemplation and research on the fascinating yet underappreciated diel ecology of insect biodiversity.

## Material and methods

### Data compilation

Relevant studies which systematically sampled insect communities across the diel cycle were identified in a literature search on 28^th^ April 2022. The full details of the procedure were described in Wong and Didham (2024); a summary is provided here. The search terms used were ‘(insect) AND (community OR communities) AND (activity OR diel OR nocturnal OR diurnal OR night OR day)’. Only data from studies using sampling methods that collected active individual insects, such as movement-based interception traps (e.g. pitfall traps, sticky traps, malaise traps, drift nets) and some attraction-based traps (e.g. dung-baited pitfall traps) was included. Data from studies using methods which potentially collected inactive individuals (e.g. sweep-netting, beating) as well as methods for which collection efficiency or attractiveness was influenced by environmental changes across the diel cycle, such as light traps and coloured pan-traps were excluded.

Across all studies that were identified as relevant, the day was considered the period after sunrise and before sunset, while the night was the period after sunset and before sunrise. For each relevant study, I recorded the total number of species collected across all samples, the numbers of species occurring exclusively in day samples (diurnal species), night samples (nocturnal species), and in both day and night samples (cathemeral species). Besides data on community composition, I recorded the season(s) during which sampling occurred as well as the elevation and geographic coordinates of the sampled locality. I then used the coordinates and the WorldClim database (Fick et al., 2017) to obtain data for 19 climate variables (see https://www.worldclim.org/data/bioclim.html) within a 1000-m radius of the locality. I also obtained the mean value of net primary productivity in the same area from the MODIS database (Justice et al., 2002) and the mean value of the Human Footprint Index (Venter et al., 2016).

### Data analysis

The proportions of diurnal, nocturnal, and cathemeral species in each community were calculated by dividing the number of species in each category over the total number of species collected across all samples. Due to the heterogenous nature of the data in study design and the absence of sample-based estimates of species richness, a formal statistical analysis was not conducted. Instead, the potential influence of locality-specific geographic and environmental factors on the proportions of diurnal, nocturnal, and cathemeral species in communities was explored by calculating summary statistics (means and SDs) and pairwise Pearson correlation coefficients, and visualising the relationships in scatterplots and bar charts. All data visualisation and analyses were performed in R software version 4.3.0 (R Core Team, 2024).

## Results

Data on the diel partitioning of species richness in 60 insect communities from 50 studies were compiled (Supporting information). These included multiple communities of the most diverse insect orders, Coleoptera, Diptera, and Hymenoptera (Fig. 1a). The sampled localities were distributed globally and had a relatively even span across the tropical and temperate latitudinal zones (−35.7–63.7) (Fig. 1a). Sampling effort varied considerably across the studies (e.g., >9000 dung-baits in Viljanen et al., 2010 versus 24 flight-interception traps in Chatzimanolis et al., 2004), however the different collection methods meant that sample sizes were not comparable indicators of sampling effort or sample coverage among the studies.

**Figure 1.**
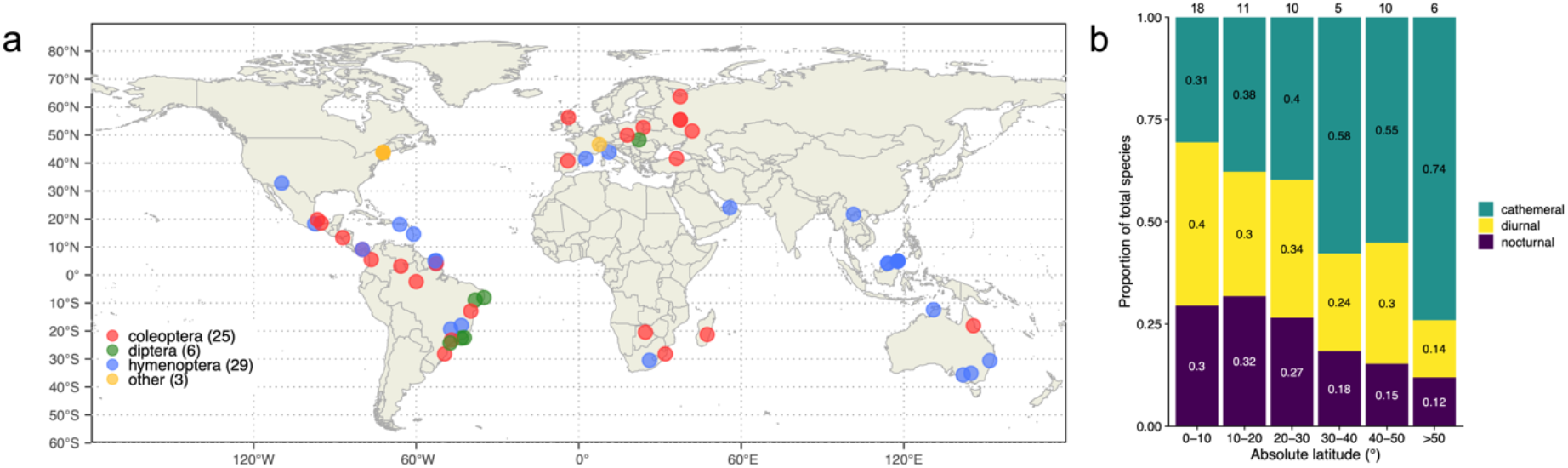
A latitudinal gradient in the diel partitioning of species richness in insect communities. **a:** The global distribution of the 60 insect communities used in the study. Numbers in parentheses indicate the number of communities included for a particular taxonomic group. **b:** The mean proportions of diurnal, nocturnal, and cathemeral species among communities within 10° latitudinal bands. The number of communities within each latitudinal band is indicated at the top its respective bar. The relative proportions of nocturnal and diurnal species tend to decrease while that of cathemeral species increases with distance from the equator.

There was high variation among the communities in total species richness (*M*=36.2, *SD*= 45.5), as well as in the relative proportions of species using the three diel activity periods. On average, diurnal species comprised 31% (*SD=*20%) of total species. The proportions of nocturnal and cathemeral species showed greater variability, averaging 24% (*SD*=21%) and 43% (*SD*=31%) of total species, respectively. The proportion of cathemeral species in a community strongly negatively correlated with the proportions of diurnal (*r*=-0.71) and nocturnal (*r*=-0.73) species, which were weaky correlated with one another (*r*=0.07).

Distinct latitudinal gradients in the diel partitioning of species richness in communities among diurnal, nocturnal, and cathemeral periods of activity were detected (Fig. 1b). The proportion of nocturnal species in a community negatively correlated with absolute latitude (*r*=-0.36) and generally declined from a mean of 30% among communities within 10° from the equator to a mean of 12% among communities beyond 50° from the equator (Fig. 1b). Similarly, the proportion of diurnal species negatively correlated with absolute latitude (*r*=-0.33) and declined poleward, from a peak of 40% within 10° from the equator to a minimum of 14% beyond 50° from the equator (Fig. 1b). In contrast, the proportion of cathemeral species positively correlated with latitude (*r*=0.45) and increased poleward from a mean of 31% within 10° from the equator, eventually doubling to 74% beyond 50° from the equator (Fig. 1b).

Besides latitude, the proportions of species using specific diel periods in communities correlated with several properties of the environmental temperature (Supporting information). Chiefly among these were two closely related indicators of the variability in temperatures over a year: temperature seasonality and the annual range in temperature. Both variables showed weak to moderate correlations (|*r*|*=*0.3–0.49) with the proportions of nocturnal, diurnal and cathemeral species in communities (Supporting information). Moreover, the directional trends of the specific relationships broadly reflected those observed with latitude: the proportions of nocturnal and diurnal species shared a similar response (i.e. a decrease) with increasing temperature seasonality and range, contrasting the proportion of cathemeral species, which increased along these climatic gradients (Supporting information). No notable correlations were observed between the proportions of species using specific diel periods and the total species richness or total abundance of insects in the communities (Supporting information).

## Discussion

The present survey of the literature on diel partitioning in insect communities provides a glimpse of the variability in these patterns globally, and reveals an interesting relationship with latitude, where poleward declines in the proportions of nocturnal and diurnal species are compensated by increments in the proportions of cathemeral species. Notably, these latitudinal trends in diel partitioning in insect communities broadly align with those observed in the richness of diurnal, nocturnal, and cathemeral mammals as well as lizards across biogeographic regions globally (Bennie et al., 2014; Vidan et al., 2019; Cox & Gaston, 2024). However, I must impress that the generality of the relationships for insects are tempered by the small sample which provides limited representation of global insect diversity, and the heterogenous data sourced from unrelated studies not originally intended for comparative analysis.

Below I outline six preliminary explanations for the apparent latitudinal gradient in the diel partitioning of species richness. The six hypotheses emphasise the roles of abiotic factors that vary with latitude, such as climate and light, biotic interactions such as competition and predation, as well as human disturbances (Table 1). This is by no means an exhaustive list; after all, no less than thirty hypotheses have been proposed to explain the classical latitudinal diversity gradient (Willig et al., 2003).

**Table 1.**
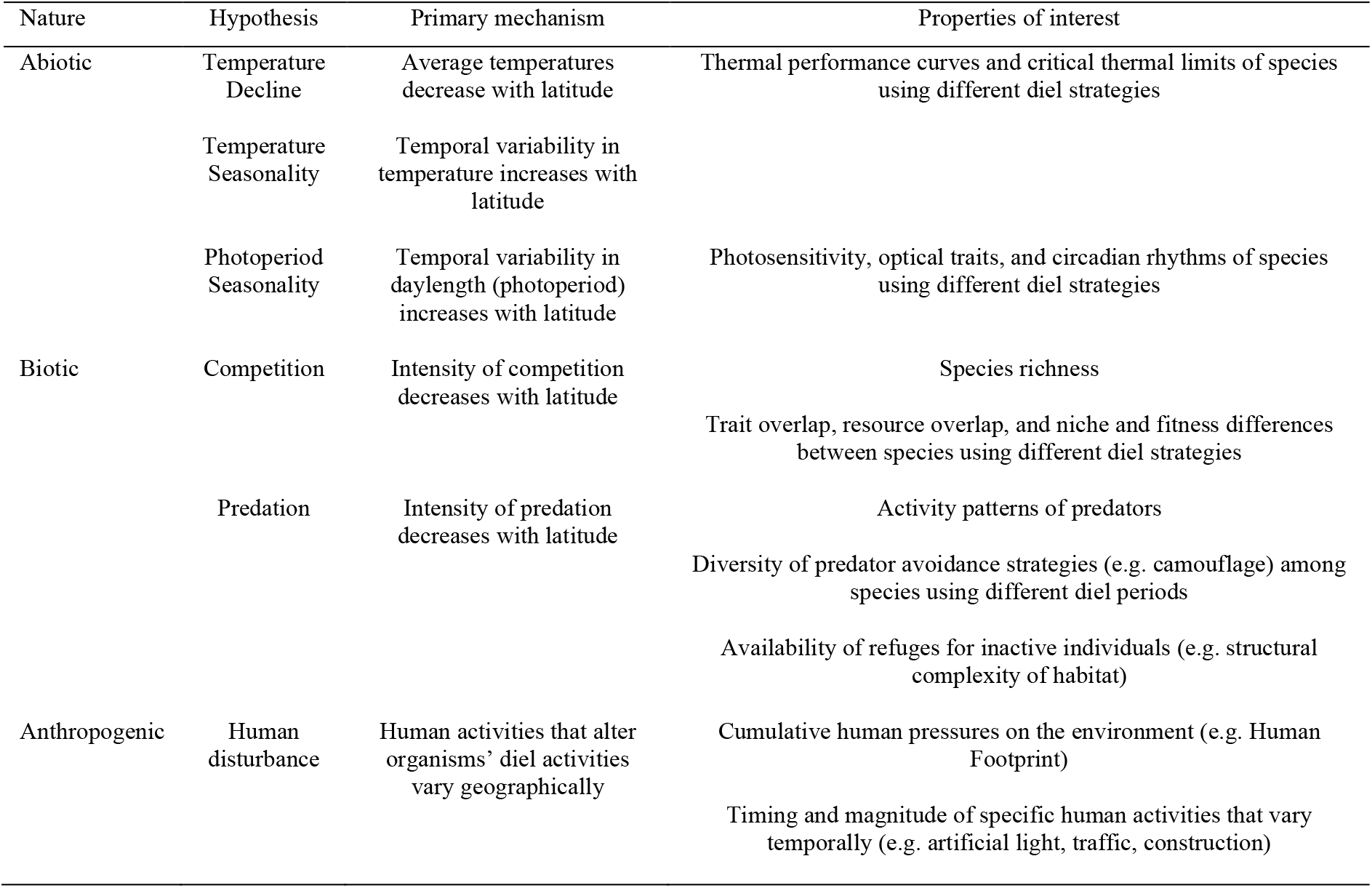
Illustration of six preliminary hypotheses that possibly account for the latitudinal gradient in the diel partitioning of species richness shown in Fig. 1, and the properties of communities and environments that may be studied to detect their underlying mechanisms.

Insects are ectotherms, making their activity levels especially constrained by environmental temperatures (Hoffmann et al., 2013). The *Temperature Decline Hypothesis* emphasises the role of the general decrease in temperature with latitude in shaping the diel partitioning of species richness in insect communities. In tropical regions, consistently high daytime temperatures may approximate species’ upper thermal limits, promoting high levels of nocturnality (Kearney et al., 2009). Conversely, nocturnality may be energetically costly in temperate regions where night temperatures approximate species’ lower thermal limits (Sinclair, 2015). To cope with these constraints, nocturnal lineages may have shifted to cathemeral activity, which is afforded by the relatively moderate daytime temperatures regionally.

The *Temperature Seasonality Hypothesis* emphasises the higher levels of seasonality and variation in temperatures that characterise environments at higher latitudes, which may limit the adaptation to strictly diurnal or nocturnal strategies (Bennie et al., 2014). In contrast, cathemeral species with comparatively flexible activity patterns may be better poised to exploit temporally variable climates for resource acquisition and reproduction. Notably, this hypothesis is supported by the similar trends in diel partitioning observed along the latitudinal gradient and the temperature seasonality gradient (i.e., decreasing proportions of nocturnal and diurnal species, increasing proportions of cathemeral species) (Supporting information).

Similarly, the *Photoperiod Seasonality Hypothesis* emphasises the pronounced seasonal variation in daylight duration (photoperiod) at high latitudes, which may exert stronger selection for a cathemeral strategy. In contrast to a strictly light-constrained strategy of nocturnal or diurnal activity, activity less constrained by a particular level of environmental light allows a species to efficiently capitalise on resource availability throughout the variable lengths of day and night in temperate regions.

Beyond abiotic factors, patterns of diel partitioning in communities may be shaped by species interactions which vary with latitude. The *Competition Hypothesis* posits that the higher species richness and more intense competition for resources in tropical communities (Pianka, 1966) promotes greater partitioning of diel activity time among species, resulting in high proportions of diurnal and nocturnal specialists, and low proportions of more generalist cathemeral species that are less well adapted to daytime or nighttime conditions. Temperate communities, which are typically characterised by lower diversity and less intense competition, may provide more opportunities for cathemeral species to exploit resources throughout the diel cycle. Interestingly, this hypothesis is not supported by the current – albeit limited – data, as the patterns of diel partitioning did not strongly correlate with total species richness (Supporting information). Nonetheless, interspecific competition has regularly been cited as key driver of diel partitioning in tropical communities of ants, bees, and dung beetles (e.g. Feer & Pincebourde, 2005; Smith et al., 2017; O’Donell et al., 2021).

The *Predation Hypothesis* emphasises the latitudinal variation in predation risk, which has been shown to peak in the tropics, especially for insects (Roslin et al., 2017). In this regard, the high diversity of nocturnal insects in tropical regions may represent a strategy to avoid diurnal predators. For instance, the richness of Lepidoptera with strictly nocturnal larvae in the Amazon has been linked to the demonstrably lower attack rates by predators during the night (Seifert et al., 2015). Notably, tropical regions often support dense vegetation and complex microhabitats, which can be exploited by nocturnal insects as refuges from diurnal predators during the day (Novotny et al., 2009). In contrast, temperate habitats may have more open and less structurally complex landscapes that afford fewer refuges for inactive nocturnal species during the day.

The *Human Disturbance Hypothesis* emphasises the role of human activities in influencing species’ activity times and patterns of diel partitioning in communities. Since human activity varies across the globe (Venter et al., 2016), it may alter biogeographic patterns in the diel partitioning of community composition. Still, such effects will largely depend on the type of human disturbances and taxa in question. For example, while many mammalian species display increased nocturnality in response to urban development (Gaynor et al., 2018), some insects show reduced nocturnal activity in the presence of artificial light (Firebaugh & Haynes, 2016). It is interesting to note that the proportions of nocturnal species in insect communities negatively correlated with the Human Footprint Index, a measure of integrated human pressures such as the amount of built infrastructure and the human population density in the landscape (Venter et al., 2016).

There is fertile ground for exploring the mechanistic basis and macroecology of the diel partitioning of community composition. Importantly, the latitudinal trends we observe (Fig. 1) are in need of hard evidence from coordinated and standardised studies of communities across regions globally (e.g. Roslin et al., 2017), focusing on diel partitioning. Complementary insights may also be acquired through similar work along elevational gradients, which mirror many aspects of environmental variation along the latitudinal gradient (Stevens, 1992). If the patterns hold true, an exciting next step will be to compare the relative contributions of different hypothetical mechanisms, such as by quantifying and modelling various mechanism-specific properties of interest (examples in Table 1).

Diel dynamics are an inherent component of biodiversity and ecosystem functioning on a rapidly changing planet. There is hence strong impetus for today’s ecologists to continue the work of the first naturalists: embracing the timeless pursuit of comprehending animal activity.

## Acknowledgements

I am indebted to the many researchers and field assistants who have worked tirelessly over the years to record crucial empirical information on the composition and activity patterns of insect communities *in situ*. A complete list of the studies from which data was gathered is found in Supporting information.

## References

Bennie, J. J., Duffy, J. P., Inger, R., & Gaston, K. J. (2014). Biogeography of time partitioning in mammals. Proceedings of the National Academy of Sciences, 111(38), 13727–13732.

Carothers, J. H., & Jaksić, F. M. (1984). Time as a niche difference: the role of interference competition. Oikos, 403–406.

Chatzimanolis, S., Ashe, J. S., & Hanley, R. S. (2004). Diurnal/nocturnal activity of rove beetles (Coleoptera: Staphylinidae) on Barro Colorado Island, Panama assayed by flight intercept trap. The Coleopterists Bulletin, 58(4), 569–577.

Cox, D. T., & Gaston, K. J. (2024). Cathemerality: a key temporal niche. Biological Reviews, 99(2), 329–347.

Darwin, C. R. 1871. The descent of man, and selection in relation to sex. London: John Murray.

Feer, F., & Pincebourde, S. (2005). Diel flight activity and ecological segregation within an assemblage of tropical forest dung and carrion beetles. Journal of Tropical Ecology, 21(1), 21–30.

Fick, S. E., & Hijmans, R. J. (2017). WorldClim 2: new 1-km spatial resolution climate surfaces for global land areas. International journal of climatology, 37(12), 4302–4315.

Firebaugh, A., & Haynes, K. J. (2016). Experimental tests of light-pollution impacts on nocturnal insect courtship and dispersal. Oecologia, 182(4), 1203–1211.

Gaston, K. J. (2019). Nighttime ecology: the “nocturnal problem” revisited. The American Naturalist, 193(4), 481–502.

Gaynor, K. M., Hojnowski, C. E., Carter, N. H., & Brashares, J. S. (2018). The influence of human disturbance on wildlife nocturnality. Science, 360(6394), 1232–1235.

Hoffmann, A. A., Chown, S. L., & Clusella-Trullas, S. (2013). Upper thermal limits in terrestrial ectotherms: how constrained are they?. Functional Ecology, 27(4), 934–949.

Kearney, M., Shine, R., & Porter, W. P. (2009). The potential for behavioral thermoregulation to buffer “cold-blooded” animals against climate warming. Proceedings of the National Academy of Sciences, 106(10), 3835–3840.

Kronfeld-Schor, N., & Dayan, T. (2003). Partitioning of time as an ecological resource. Annual review of ecology, evolution, and systematics, 34(1), 153–181.

Janzen, D. H., & Hallwachs, W. (2021). To us insectometers, it is clear that insect decline in our Costa Rican tropics is real, so let’s be kind to the survivors. Proceedings of the National Academy of Sciences, 118(2), e2002546117.

Justice, C. O., Townshend, J. R. G., Vermote, E. F., Masuoka, E., Wolfe, R. E., Saleous, N.,… & Morisette, J. T. (2002). An overview of MODIS Land data processing and product status. Remote sensing of Environment, 83(1-2), 3–15.

Lear, K. O., Whitney, N. M., Morris, J. J., & Gleiss, A. C. (2021). Temporal niche partitioning as a novel mechanism promoting co-existence of sympatric predators in marine systems. Proceedings of the Royal Society B, 288(1954), 20210816.

Miraldo, A., Li, S., Borregaard, M. K., Flórez-Rodríguez, A., Gopalakrishnan, S., Rizvanovic, M.,… & Nogués-Bravo, D. (2016). An Anthropocene map of genetic diversity. Science, 353(6307), 1532–1535.

Novotny, V., Basset, Y., Auga, J., Boen, W., Dal, C., Drozd, P.,… & Manumbor, M. (1999). Predation risk for herbivorous insects on tropical vegetation: a search for enemy-free space and time. Australian Journal of Ecology, 24(5), 477–483.

O’Donnell, S., Lattke, J., Powell, S., & Kaspari, M. (2021). Diurnal and nocturnal foraging specialisation in Neotropical army ants. Ecological Entomology, 46(2).

Otté E. C. & Bohn, H. B. (1850). Views of Nature: or Contemplations on the Sublime Phenomena of Creation with Scientific Illustrations, 3rd edn (Alexander von Humboldt), Henry G. Bohn.

Pianka, E. R. (1966). Latitudinal gradients in species diversity: a review of concepts. The American Naturalist, 100(910), 33–46.

R Core Team. (2024). A language and environment for statistical computing.

Roslin, T., Hardwick, B., Novotny, V., Petry, W. K., Andrew, N. R., Asmus, A.,… & Slade, E. M. (2017). Higher predation risk for insect prey at low latitudes and elevations. Science, 356(6339), 742–744.

Sanders, D., Frago, E., Kehoe, R., Patterson, C., & Gaston, K. J. (2021). A meta-analysis of biological impacts of artificial light at night. Nature Ecology & Evolution, 5(1), 74–81.

Schumm, M., Edie, S. M., Collins, K. S., Gómez-Bahamón, V., Supriya, K., White, A. E.,… & Jablonski, D. (2019). Common latitudinal gradients in functional richness and functional evenness across marine and terrestrial systems. Proceedings of the Royal Society B, 286(1908), 20190745.

Seifert, C. L., Schulze, C. H., Dreschke, T. C., Frötscher, H., & Fiedler, K. (2016). Day vs. night predation on artificial caterpillars in primary rainforest habitats–an experimental approach. Entomologia Experimentalis et Applicata, 158(1), 54–59.

Sinclair, B. J. (2015). Linking energetics and overwintering in temperate insects. Journal of Thermal Biology, 54, 5–11.

Smith, A. R., Kitchen, S. M., Toney, R. M., & Ziegler, C. (2017). Is nocturnal foraging in a tropical bee an escape from interference competition?. Journal of Insect Science, 17(2), 62.

Stevens, G. C. (1992). The elevational gradient in altitudinal range: an extension of Rapoport’s latitudinal rule to altitude. The American Naturalist, 140(6), 893–911.

Tattersall, I. (1987). Cathemeral activity in primates: a definition. Folia Primatol, 49(3-4), 200–202.

Thoré, E. S., Aulsebrook, A. E., Brand, J. A., Almeida, R. A., Brodin, T., & Bertram, M. G. (2024). Time is of the essence: The importance of considering biological rhythms in an increasingly polluted world. Plos Biology, 22(1), e3002478.

Venter, O., Sanderson, E. W., Magrach, A., Allan, J. R., Beher, J., Jones, K. R.,… & Watson, J. E. (2016). Global terrestrial Human Footprint maps for 1993 and 2009. Scientific data, 3(1), 1–10.

Vidan, E., Novosolov, M., Bauer, A. M., Herrera, F. C., Chirio, L., de Campos Nogueira, C.,… & Meiri, S. (2019). The global biogeography of lizard functional groups. Journal of Biogeography, 46(10), 2147–2158.

Viljanen, H., Wirta, H., Montreuil, O., Rahagalala, P., Johnson, S., & Hanski, I. (2010). Structure of local communities of endemic dung beetles in Madagascar. Journal of Tropical Ecology, 26(5), 481–496.

Willig, M. R., Kaufman, D. M., & Stevens, R. D. (2003). Latitudinal gradients of biodiversity: pattern, process, scale, and synthesis. Annual review of ecology, evolution, and systematics, 34(1), 273–309.

Wong, M. K. L., & Didham, R. K. (2024). Global meta-analysis reveals overall higher nocturnal than diurnal activity in insect communities. Nature Communications, 15(1), 3236.

