## Supporting information for "A latitudinal gradient in the diel partitioning of species richness?"

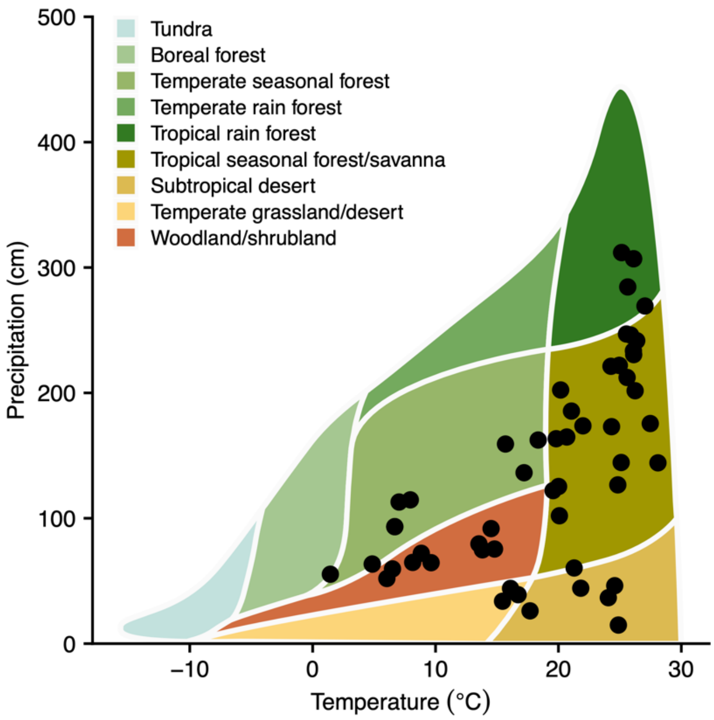


**Figure S1.** Distribution of the 60 insect communities across climate space as encapsulated by Whittaker's biomes.

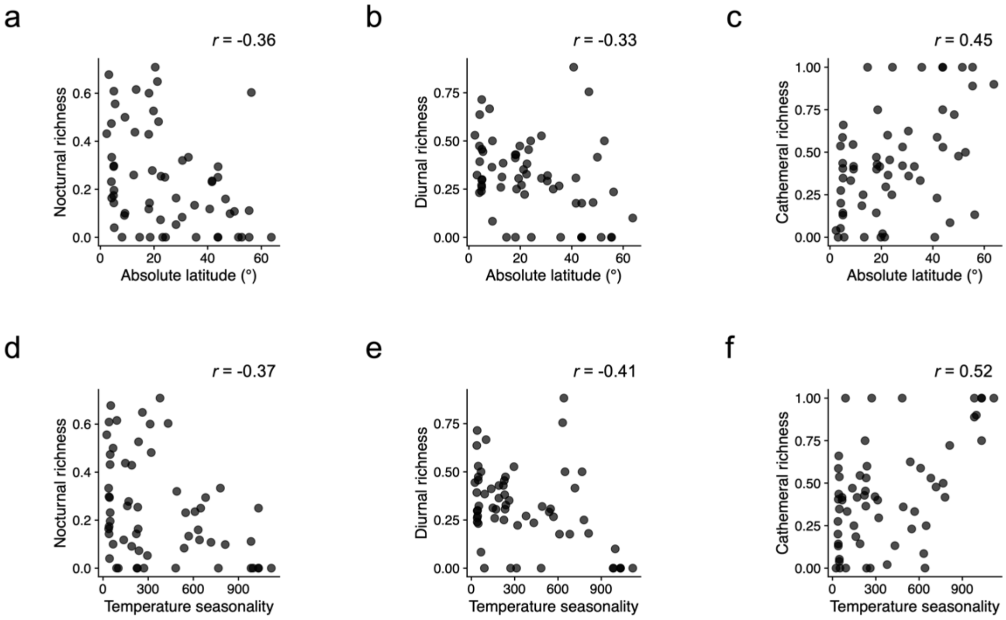


**Figure S2.** Scatterplots of the proportions of nocturnal, diurnal, and cathemeral species in 60 insect communities along gradients of absolute latitude (a–c) and temperature seasonality (d–f). Correlation coefficient values (Pearson's *r*) are shown on the top right of each plot. Temperature seasonality is a measure of the annual variation in atmospheric temperatures at a locality. Data on temperature seasonality was obtained from WorldClim (bioclimatic variable 'BIO14').

**Table S1.** Summary statistics for the values of total species richness and the relative proportions of diurnal, nocturnal, and cathemeral species in the 60 studied insect communities.

| Group | N | Total species | | Diurnal | | Nocturnal | | Cathemeral | |
|---|---|---|---|---|---|---|---|---|---|
| | | *M* | *SD* | *M* | *SD* | *M* | *SD* | *M* | *SD* |
| Combined | 60 | 36.22 | 45.53 | 0.31 | 0.20 | 0.24 | 0.21 | 0.43 | 0.31 |
| Coleoptera | 25 | 27.48 | 19.23 | 0.32 | 0.22 | 0.30 | 0.26 | 0.38 | 0.35 |
| Diptera | 6 | 35.67 | 27.02 | 0.32 | 0.22 | 0.09 | 0.09 | 0.59 | 0.25 |
| Hymenoptera | 26 | 36.15 | 29.43 | 0.31 | 0.16 | 0.24 | 0.15 | 0.42 | 0.23 |
| Other Insecta | 3 | 110.67 | 186.48 | 0.25 | 0.44 | 0.05 | 0.09 | 0.70 | 0.53 |

**Table S2.** Pairwise Pearson correlation coefficients (*r*) between individual biotic, geographical, climatic and disturbance-related properties of a locality and the relative proportions of nocturnal, diurnal and cathemeral species in the respective insect community (n=60). Coefficients indicating weak to moderate associations ($|r|\geq0.3$) are highlighted in bold.

| | | **Nocturnal richness** | **Diurnal richness** | **Cathemeral richness** |
|---|---|---|---|---|
| *Biota* | | | | |
| | Total richness | 0.05 | **0.36** | -0.25 |
| | Total abundance | -0.06 | -0.20 | 0.17 |
| | Net primary productivity (mean) | 0.05 | 0.20 | -0.15 |
| *Geography* | | | | |
| | Latitude (absolute) | **-0.36** | **-0.33** | **0.45** |
| | Elevation | 0.09 | 0.34 | -0.26 |
| *Climate* | | | | |
| | Temperature (mean) | **0.38** | 0.29 | **-0.46** |
| | Temperature (maximum) | 0.27 | 0.08 | -0.26 |
| | Temperature (minimum) | **0.36** | **0.36** | **-0.48** |
| | Temperature (isothermality) | **0.36** | **0.35** | **-0.46** |
| | Temperature (seasonality) | **-0.37** | **-0.41** | **0.52** |
| | Temperature (annual range) | **-0.30** | **-0.40** | **0.46** |
| | Precipitation (mean) | **0.30** | 0.20 | **-0.32** |
| | Precipitation (maximum) | **0.36** | 0.17 | **-0.36** |
| | Precipitation (minimum) | 0.18 | 0.18 | -0.21 |
| | Precipitation (seasonality) | **0.30** | 0.12 | **-0.30** |
| *Human impact* | | | | |
| | Human footprint index (mean) | **-0.37** | -0.05 | 0.26 |
| | Artificial sky luminance (mean) | -0.29 | 0.00 | 0.18 |

**Supplementary Note 1.** Reference list of studies on insect communities from which data on the diel partitioning species richness was obtained.


1. Alencar, J., Fereira, Z. M., Lopes, C. M., Serra-Freire, N. M., De Mello, R. P., Silva, J. D. S., & Guimarães, A. É. (2011). Biodiversity and times of activity of mosquitoes (Diptera: Culicidae) in the biome of the Atlantic Forest in the State of Rio de Janeiro, Brazil. Journal of medical entomology, 48(2), 223-231.
2. Amézquita, S., & Favila, M. E. (2011). Carrion removal rates and diel activity of necrophagous beetles (Coleoptera: Scarabaeinae) in a fragmented tropical rain forest. Environmental Entomology, 40(2), 239-246.
3. Andersen, A. N. (1983). Species diversity and temporal distribution of ants in the semi_arid mallee region of northwestern Victoria. Australian Journal of Ecology, 8(2), 127-137.
4. Andresen, E. (2002). Dung beetles in a Central Amazonian rainforest and their ecological role as secondary seed dispersers. Ecological Entomology, 27(3), 257-270.
5. Anjos, D. V., Caserio, B., Rezende, F. T., Ribeiro, S. P., Del_Claro, K., & Fagundes, R. (2017). Extrafloral_nectaries and interspecific aggressiveness regulate day/night turnover of ant species foraging for nectar on Bionia coriacea. Austral Ecology, 42(3), 317-328.
6. Briese, D. T., & Macauley, B. J. (1980). Temporal structure of an ant community in semi_arid Australia. Australian Journal of Ecology, 5(2), 121-134.
7. Carval, D., Cotté, V., Resmond, R., Perrin, B., & Tixier, P. (2016). Dominance in a ground_dwelling ant community of banana agroecosystem. Ecology and Evolution, 6(23), 8617-8631.
8. Cerdá, X., Retana, J., & Cros, S. (1998). Critical thermal limits in Mediterranean ant species: trade_off between mortality risk and foraging performance. Functional Ecology, 12(1), 45-55.
9. Chatzimanolis, S., Ashe, J. S., & Hanley, R. S. (2004). Diurnal/nocturnal activity of rove beetles (Coleoptera: Staphylinidae) on Barro Colorado Island, Panama assayed by flight intercept trap. The Coleopterists Bulletin, 58(4), 569-577.
10. Chew, R. M. (1977). Some ecological characteristics of the ants of a desert-shrub community in southeastern Arizona. American Midland Naturalist, 33-49.
11. Cole, L. J., McCracken, D. I., Dennis, P., Downie, I. S., Griffin, A. L., Foster, G. N., ... & Waterhouse, T. (2002). Relationships between agricultural management and ecological groups of ground beetles (Coleoptera: Carabidae) on Scottish farmland. Agriculture, Ecosystems & Environment, 93(1-3), 323-336.
12. da Silva, P. G., Lobo, J. M., & Hernández, M. I. M. (2019). The role of habitat and daily activity patterns in explaining the diversity of mountain Neotropical dung beetle assemblages. Austral Ecology, 44(2), 300-312.
13. de Oca T, E. M., & Halffter, G. (1995). Daily and seasonal activities of a guild of the coprophagous, burrowing beetle (Coleoptera Scarabaeidae Scarabaeinae) in tropical grassland. Tropical Zoology, 8(1), 159-180.
14. Doube, B. M. (1983). The habitat preference of some bovine dung beetles (Coleoptera: Scarabaeidae) in Hluhluwe Game Reserve, South Africa. Bulletin of Entomological Research, 73(3), 357-371.

15. Feer, F., & Pincebourde, S. (2005). Diel flight activity and ecological segregation within an assemblage of tropical forest dung and carrion beetles. Journal of Tropical Ecology, 21(1), 21-30.
16. Forattini, O. P., Alves, A. D. C., Natal, D., & Santos, J. L. F. (1986). Observações sobre atividade de mosquitos Culicidae em mata primitiva da encosta no Vale do Ribeira, São Paulo, Brasil. Revista de Saúde Pública, 20, 1-20.
17. Frizzi, F., Tucci, L., Ottonetti, L., Masoni, A., & Santini, G. (2021). Day-night and inter-habitat variations in ant assemblages in a mosaic agroforestry landscape. Land, 10(2), 179.
18. Grevé, M. E., Houadria, M., Andersen, A. N., & Menzel, F. (2019). Niche differentiation in rainforest ant communities across three continents. Ecology and Evolution, 9(15), 8601-8615.
19. Gryuntal, S. Y., & Ã__íòàëü, Ñ. _. (2008). Daily activity of carabid beetles (Coleoptera: Carabidae) in the forests of various geographical zones in the East European (Russian) plain. Russian Entomological Journal, 17, 359-365.
20. Guimarães, A. É., Gentile, C., Lopes, C. M., & Mello, R. P. D. (2000). Ecology of mosquitoes (Diptera: Culicidae) in areas of Serra do Mar State Park, State of São Paulo, Brazil. III-daily biting rhythms and lunar cycle influence. Memórias do Instituto Oswaldo Cruz, 95, 753-760.
21. Hampton, S. E., & Friedenberg, N. A. (2002). Nocturnal increases in the use of near-surface water by pond animals. Hydrobiologia, 477, 171-179.
22. Hashimoto, Y., Morimoto, Y., Widodo, E. S., Mohamed, M., & Fellowes, J. R. (2010). Vertical habitat use and foraging activities of arboreal and ground ants (Hymenoptera: Formicidae) in a Bornean tropical rainforest. Sociobiology, 56(2), 435.
23. Heatwole, H. (1991). The ant assemblage of a sand-dune desert in the United Arab Emirates. Journal of arid environments, 21(1), 71-79.
24. Heatwole, H., Trémont, S., & Broese, E. (2013). Point-diversity, a critical tool for assessing dynamics of guilds of scavenging ants (Hymenoptera: Formicidae): an example from a eucalypt woodland. Systematics and biodiversity, 11(2), 149-180.
25. Hernández, M. I. M. (2002). The night and day of dung beetles (Coleoptera, Scarabaeidae) in the Serra do Japi, Brazil: elytra colour related to daily activity. Revista brasileira de Entomologia, 46, 597-600.
26. Hill, C. J. (1996). Habitat specificity and food preferences of an assemblage of tropical Australian dung beetles. Journal of Tropical Ecology, 12(4), 449-460.
27. Horgan, F. G. (2002). Shady field boundaries and the colonisation of dung by coprophagous beetles in Central American pastures. Agriculture, ecosystems & environment, 91(1-3), 25-36.
28. Hossaert-McKey, M., Orivel, J., Labeyrie, E., Pascal, L., Delabie, J., & Dejean, A. (2001). Differential associations with ants of three cooccurring extrafloral nectary-bearing plants. Ecoscience, 8(3), 325-335.
29. Kami_ski, M. J., Byk, A., & Tykarski, P. (2015). Seasonal and diel activity of dung beetles (Coleoptera: Scarabaeoidea) attracted to European bison dung in Bia_owie_a Primeval Forest, Poland. The Coleopterists Bulletin, 69(1), 83-90.
30. Kaspari, M., & Weiser, M. D. (2000). Ant activity along moisture gradients in a neotropical forest 1. Biotropica, 32(4a), 703-711.

31. Kirmse, S., & Ratcliffe, B. C. (2019). Composition and host-use patterns of a scarab beetle (Coleoptera: Scarabaeidae) community inhabiting the canopy of a lowland tropical rainforest in southern Venezuela. The Coleopterists Bulletin, 73(1), 149-167.
32. Knop, E., Gerpe, C., Ryser, R., Hofmann, F., Menz, M. H., Trösch, S., ... & Fontaine, C. (2018). Rush hours in flower visitors over a day–night cycle. Insect Conservation and Diversity, 11(3), 267-275.
33. Ko_árek, P. (2002). Diel activity patterns of carrion-visiting Coleoptera studied by time-sorting pitfall traps.
34. Lindsey, P. A., & Skinner, J. D. (2001). Ant composition and activity patterns as determined by pitfall trapping and other methods in three habitats in the semi-arid Karoo. Journal of Arid Environments, 48(4), 551-568.
35. Liu, X., Wang, Z., Huang, C., Li, M., Bibi, F., Zhou, S., & Nakamura, A. (2020). Ant assemblage composition explains high predation pressure on artificial caterpillars during early night. Ecological Entomology, 45(3), 547-554.
36. Lobo, J. M., & Cuesta, E. (2021). Seasonal variation in the diel activity of a dung beetle assemblage. PeerJ, 9, e11786.
37. Luna, P., Peñaloza-Arellanes, Y., Castillo-Meza, A. L., García-Chávez, J. H., & Dáttilo, W. (2018). Beta diversity of ant-plant interactions over day-night periods and plant physiognomies in a semiarid environment. Journal of arid environments, 156, 69-76.
38. Marques, G. D. V., & Del-Claro, K. (2006). The ant fauna in a Cerrado area: the influence of vegetation structure and seasonality (Hymenoptera: Formicidae). Sociobiology, 47(1), 235-252.
39. Medina, A. M., & Lopes, P. P. (2014). Resource utilization and temporal segregation of Scarabaeinae (Coleoptera, Scarabaeidae) community in a Caatinga fragment. Neotropical entomology, 43, 127-133.
40. Móra, A., Dévai, G., Tóthmérész, B., & Csépes, E. (2006). Short-time changes in composition of chironomid assemblages at a cross-section of the River Tisza. Internationale Vereinigung für theoretische und angewandte Limnologie: Verhandlungen, 29(4), 2099-2102.
41. Oliveira, D. L., & Vasconcelos, S. D. (2018). Diversity, daily flight activity and temporal occurrence of necrophagous Diptera associated with decomposing carcasses in a semi-arid environment. Neotropical entomology, 47, 470-477.
42. Pardo-Locarno, L. C. (2007). Escarabajos Coprófagos (coleoptera-scarabaeidae) de Lloró, departamento del Chocó, Colombia. Boletín Científico. Centro de Museos. Museo de Historia Natural, 11(1), 377-388.
43. Sands, B., Mgidiswa, N., Curson, S., Nyamukondiwa, C., & Wall, R. (2022). Dung beetle community assemblages in a southern African landscape: niche overlap between domestic and wild herbivore dung. Bulletin of Entomological Research, 112(1), 131-142.
44. Soares, T. F., & Vasconcelos, S. D. (2016). Diurnal and nocturnal flight activity of blow flies (Diptera: Calliphoridae) in a rainforest fragment in Brazil: implications for the colonization of homicide victims. Journal of forensic sciences, 61(6), 1571-1577.
45. Sullivan, G. T., Ozman-Sullivan, S. K., Lumaret, J. P., Bourne, A., Zeybekoglu, U., Zalucki, M. P., & Baxter, G. (2017). How guilds build success; aspects of temporal resource partitioning in a warm,

temperate climate assemblage of dung beetles (Coleoptera: Scarabaeidae). Environmental entomology, 46(5), 1060-1069.

46. Tanaka, H. O., Yamane, S., & Itioka, T. (2010). Within-tree distribution of nest sites and foraging areas of ants on canopy trees in a tropical rainforest in Borneo. Population Ecology, 52, 147-157.
47. Torres, J. A. (1984). Niches and coexistence of ant communities in Puerto Rico: repeated patterns. Biotropica, 284-295.
48. Viljanen, H., Wirta, H., Montreuil, O., Rahagalala, P., Johnson, S., & Hanski, I. (2010). Structure of local communities of endemic dung beetles in Madagascar. Journal of Tropical Ecology, 26(5), 481-496.
49. Yamane, S. K., Itino, T., & Nona, A. R. (1996). Ground ant fauna in a Bornean dipterocarp forest. Raffles Bulletin of Zoology, 44(1), 253-262.
50. Yusah, K. M., Foster, W. A., Reynolds, G., & Fayle, T. M. (2018). Ant mosaics in Bornean primary rain forest high canopy depend on spatial scale, time of day, and sampling method. PeerJ, 6, e4231.
